# The puzzling case of the 3p25 homozygous deletion: the target is not *VHL* but *VGLL4* and *ATG7*

**DOI:** 10.1101/2023.11.06.565014

**Authors:** Florian Muller, Eliot Behr

## Abstract

The 3p25 locus experiences recurrent homozygous deletions in diverse carcinomas. A validated tumor suppressor gene (TSG), *VHL* has long been assumed to be the target of this deletion.

Here, we show that homozygous deletions at the 3p25 locus are common in renal clear cell carcinoma (RCC) and squamous cell carcinomas but that these do not center on *VHL* but on two nearby overlapping genes, *VGLL4* and *ATG7*. While hypomorphic mutations of *VHL* coupled with heterozygous deletions are undoubtedly a driver in RCC, DepMap CRISPR data indicate that the complete absence of *VHL* is broadly detrimental to cancer cells. Re-examination of TCGA and PDX data calls into question whether *VHL* is ever homozygous deleted, even in RCC.

Instead, multiple lines of evidence support the homozygous deletion of *VGLL4* as a driver event in squamous cell carcinomas and the homozygous deletion of *ATG7* as a driver in RCC, with important implication for precision oncology. VGLL4 is the major negative regulator of the YAP/TEAD complex, with elimination of VGLL4 leading to hyperactive oncogenic YAP-dependent transcriptional activity which could be targeted by clinically emerging YAP/TEAD inhibitors. ATG7 is a critical regulator of autophagy, limiting proliferation in conditions of nutrient limitation and its deletion may open novel vulnerabilities for synthetic lethality.

## Introduction

The long arm of Chromosome 3 (3p) experiences frequent loss of heterozygosity (LOH – or monoallelic deletion) in diverse cancer types (Huebner, 2001) (Braga et al., 2002) which is especially common in RCC (Hsieh et al., 2018) and squamous cell carcinomas of diverse tissues of origin and associated with poor prognosis (Kim et al., 2021)(Taylor et al., 2018). Chromosome 3p is well known for several two-hit – heterozygous deletion + LOF mutation type-tumor suppressor genes such as *BAP1* (Han et al., 2021) and *VHL* (Hsieh et al., 2018). Along with VHL, located on the 3p25 locus specifically, there is also in vitro evidence of the 3p25 gene *XPC* as a potential tumor-suppressor (Cui et al., 2015).

Yet it is not widely appreciated that the 3p25 region also experiences recurrent highly focal homozygous (bi-allelic) deletions (Iijima et al., 2004) especially in squamous cell carcinomas (SCC). For example, ∼5% of Lung squamous cell carcinomas have homozygous deletions of the 3p25 locus. The 3p25 homozygous deletion seems to have been overlooked in TCGA as no target TSG or minimal common region of the 3p25 homozygous deletion was described in any of the major TCGA consortium papers discussed even in cancer types where it is quite common like Lung and Esophageal SCC (Cancer Genome Atlas Research Network, 2012) (Cancer Genome Atlas Research Network et al., 2017). No mention of 3p25 homozygous deletions were made in the RCC TCGA consortium paper either (Cancer Genome Atlas Research Network, 2013).

Here, we undertook a comprehensive analysis of TCGA and CrownBio PDX genomic and transcriptomic data of the 3p25 region. We show that homozygous deletions recurrently occur in this region across multiple tumor types but that these are not centered on the obvious TSG candidate – *VHL* – even in RCC. A careful review of the segmentation TCGA and CrownBio PDX data suggests that homozygous deletions of *VHL* are either rare or do not occur at all even in RCC. DepMap data show that CRISPR ablation of *VHL* is strongly detrimental and reduces fitness even in *VHL* mutant RCC cell lines. These data indicate that while deficiency in *VHL* drives oncogenesis – complete extinction is detrimental.

Instead – our analysis reveals heretofore undescribed recurrent homozygous deletions centered on *VGLL4* and *ATG7* – two genes on opposite strands with partially overlapping exons. CRISPR and mouse GEMM data indicate that loss of both *ATG7* and *VGLL4* likely represent driver events which for *ATG7* is further reinforced by Mendelian cholangiocarcinoma with mutated *ATG7* where the remain intact allele is lost by chromosome arm deletion (Greer et al., 2022). Although *ATG7* and *VGLL4* are understood to be tumor suppressor genes, even high-profile dedicated publications appear unaware that *ATG7* and *VGLL4* are in a locus of recurrent homozygous deletion {e.g. (Yamaguchi, 2020)(Mandelbaum et al., 2015)(Greer et al., 2022)(Lee et al., 2012)(Long et al., 2022)}.

Our findings have important implications for precision oncology – identifying *VGLL4* and *ATG7* homozygous deletions as potentially targetable vulnerabilities and paradoxically highlighting *VHL* as a potential oncology drug target in cancers outside of RCC.

## Methods

Key bioinformatic databases queried in the present investigation (**Table I**) are:

- cBioPortal (cbioportal.org) for TCGA tumor RNA sequencing, WES point mutation and SNP 6.0 copy number data (de Bruijn et al., 2023)
- CrownBio HuBase^TM^ (hubase.crownbio.com/PDXmodel/HuPrime) for PDX Exome seq and copy number data (Qian et al., 2022)
- DepMap (depmap.org/portal) for CRISPR viability data in cancer cell line (Barretina et al., 2012)
- TCGA Copy Number Portal (portals.broadinstitute.org/tcga/home/570730/) for GISTIC focality and recurrence analysis of TCGA SNP 6.0 data (Mermel et al., 2011)
- MGI (www.informatics.jax.org/) for knockout mouse phenotypes

**Table I.**
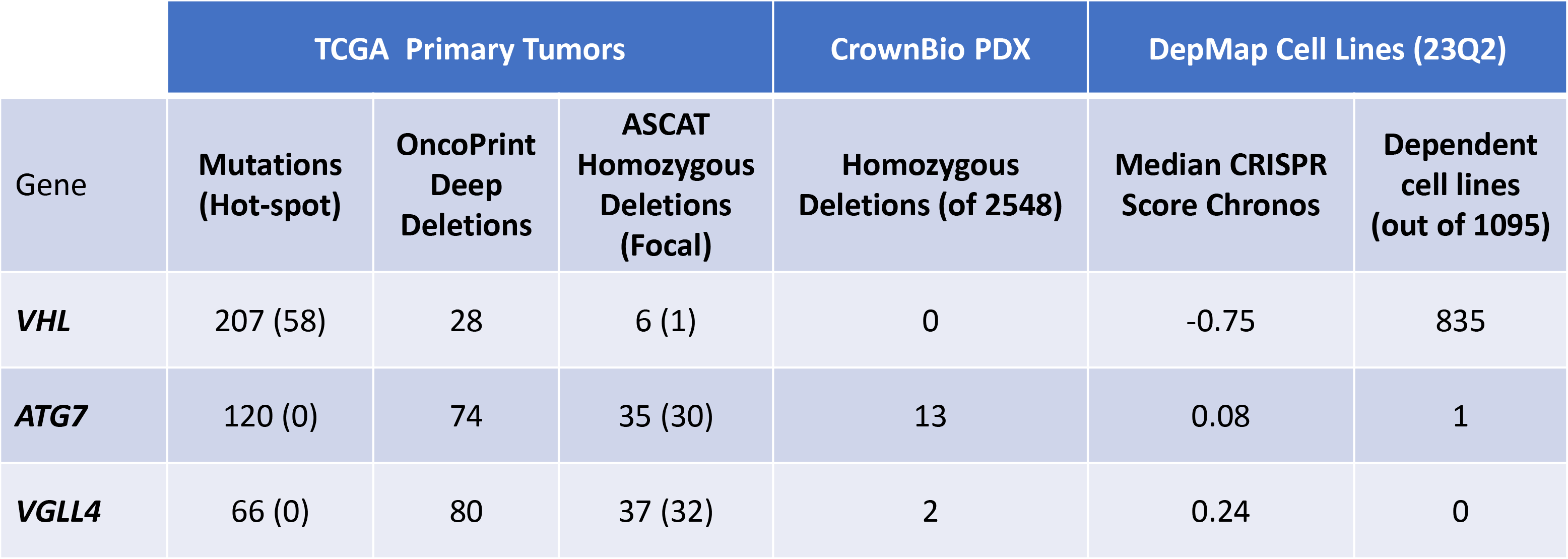

|  | TCGA Primary Tumors |  |  | CrownBio PDX | DepMap Cell Lines (23Q2) |  |
| --- | --- | --- | --- | --- | --- | --- |
| Gene | Mutations (Hot-spot) | OncoPrint Deep Deletions | ASCAT Homozygous Deletions (Focal) | Homozygous Deletions (of 2548) | Median CRISPR Score Chronos | Dependent cell lines (out of 1095) |
| *VHL* | 207 (58) | 28 | 6 (1) | 0 | -0.75 | 835 |
| *ATG7* | 120 (0) | 74 | 35 (30) | 13 | 0.08 | 1 |
| *VGLL4* | 66 (0) | 80 | 37 (32) | 2 | 0.24 | 0 |

## Results

### Challenges for reliable identification of homozygous deletions in bulk tumor resections

Homozygous deletions of the 3p25 locus were investigated in the segmented SNP 6.0 copy number variation (CNV) dataset of TCGA tumors with ∼10,000 cases across a broad range of tumor types (**Figure 1**) using cBioPortal. As precautionary note – the faithful identification homozygous deletions in data generated from bulk resected or biopsied tumors is complicated by variable admixture of genomically intact non-malignant stromal cells as well as whole genome duplications and variable ploidy in the malignant cancer cells themselves (Cheng et al., 2017). OncoPrint cBioportal data “homozygous deletion” calls on TCGA SNP 6.0 data must be interpreted with great caution since no correction for the abovementioned complications has been made. To illustrate this point: even genes that are unquestionably indispensable for cancer cell viability such as *CDK1*; *PCNA*; *POLA2* (**Figure 2**) are implausibly labeled “HOMODELETED” (homozygous deleted) in OncoPrint cBioportal summaries of TCGA SNP6.0 data (**Figure 3A**). These “HOMODELETED” calls most likely indicate heterozygous deletions and indicate that the cBioPortal OncoPrint “HOMODELETED” calls must be interpreted with caution and independently corroborated.

**Figure 1:**
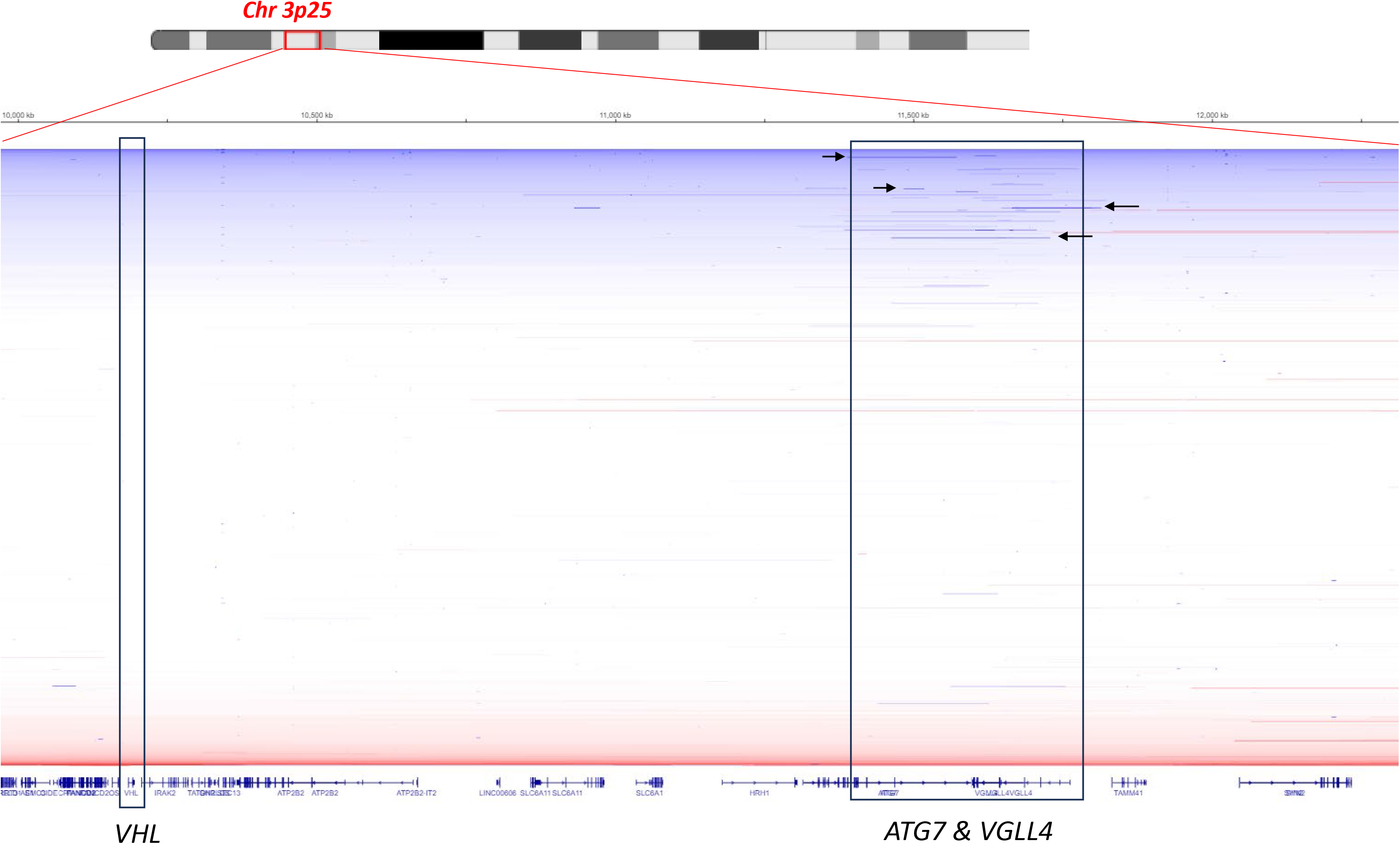
Recurrent homozygous deletions in 3p25 region center on *VGLL4* and *ATG7* but not *VHL*. Segmented normalized Log2 transformed array comparative hybridization (cGH) data SNP 6.0 data from TCGA tumors (∼10,000 cases) retrieved via cBioPortal and visualized with IGV – with chromosomal position and mapped genes (hg19) in the x-axis. Blue represents copy number loss with intensity indicating lower copy number; white neutral; red copy number gain. Homozygous deletions are evident as focal bands with lower copy number (darker blue) represent regions of homozygous deletion. Examples of homozygous deletions in the *ATG7/VGLL4* locus are indicated by black arrows. None are evident in the *VHL* locus.

**Figure 2:**
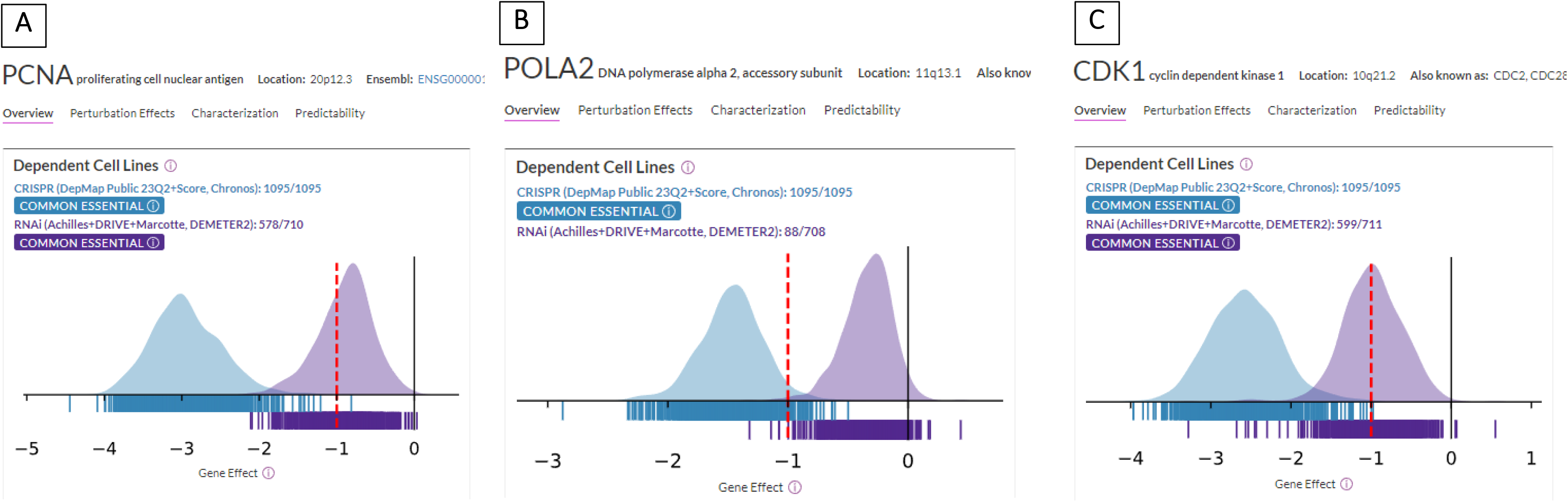
*PCNA*, *POLA2* and *CDK1* are essential in all DepMap cancer cell lines. DepMap (23Q2) data were interrogated for CRISPR Dependency scores for genes known to play non-redundant roles in cell replication. The Broad’s DepMap portal (https://depmap.org/portal/) was interrogated for CRISPR and RNAi dependencies of genes with obvious critical cellular function – replication and mitosis. Summary of CRISPR Chronos scores for 23Q2 data are shown for *PCNA* (**Panel A**), *POLA2* (**Panel B**) and *CDK1* (**Panel C**) – negative values indicate that CRISPR ablation is detrimental whilst positive values indicate that CRISPR ablation increases fitness and stimulates cancer cell proliferation. Chronos values below −1 indicate essential genes which is indicated by a red dashed line. Each bar represents one cell line. Summary data for CRISPR dependency scores for *PCNA*, *POLA2* and *CDK1* indicate that these genes are essential in 1095/1095 cancer cell lines – fully consistent with their role in [cancer] cell replication.

**Figure 3:**
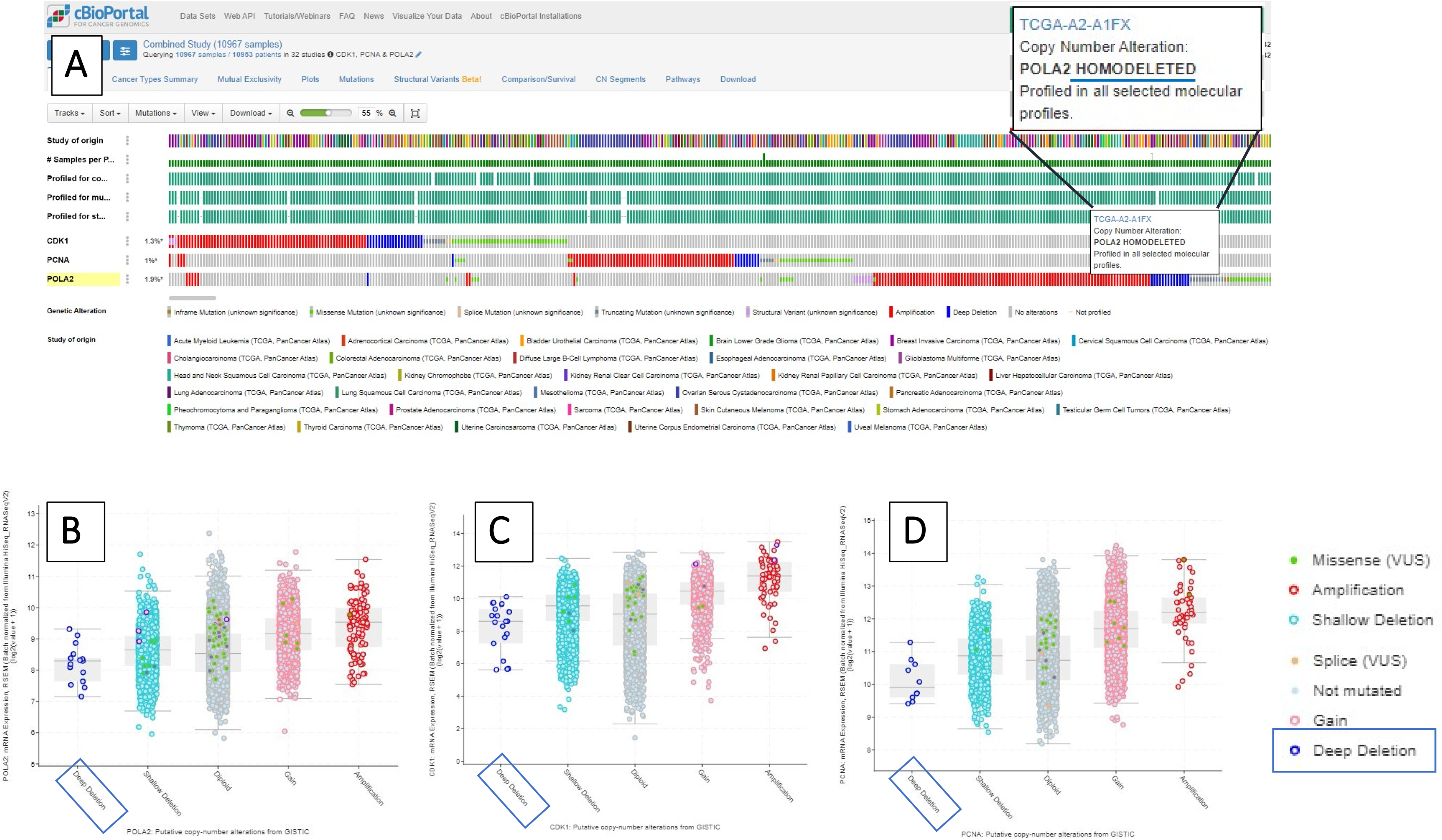
Improbable OncoPrint cBioPortal Homozygous Deletion calls in genes indispensable for cell viability. OncoPrint results of the queries in cBioPortal for the core essential genes *CDK1, PCNA* and *POLA2* in the pan-TCGA data set are shown in Panel A. Cases color coded in blue are identified as “HOMODELETED” (Homozygous) with an example for a case with *POLA2* “HOMODELETED” highlighted and magnified. Bar graphs of CNV versus mRNA expression are shown for the same genes in Panels B, C and D but with the cases identified as “HOMODELETED” in the OncoPrint now referred to as “Deep Deletion” in these plots. Note the modest decreases in mRNA expression in the cases with “Deep Deletion” which is inconsistent with the highly improbably claim of homozygous deletion of genes critical for cancer cell viability. This is in contrast with the dramatic (several Log2-fold) decreases in mRNA in cases with plausible homozygous deletion of *VGLL4* and *ATG7*.

For other cBioPortal presentation of the CNV data – for example in CNV versus mRNA expression data (**Figure 3B**) - the term “Deep deletion” rather than “HOMODELETED” is used for tumors with low CNV scores. The term “Deep deletion” more correctly reflects the nuances and difficulty of identifying actual homozygous deletions versus heterozygous deletions in the context of whole genome duplication and admixed tumors. A reliable but rather tedious method of verifying homozygous deletions consists of manual inspection of individual tumors segmentation data in IGV – where abrupt and focal decrease in Log2 copy number readily distinguish homozygous from heterozygous deletions (**Figure 1**). Other methods of triangulation consist of quantification of non-malignant stromal admixture through mRNA expression and correction for ploidy (Cheng et al., 2017) – available at COSMIC database at the Sanger Center – **Table I**). Finally, analysis of patient-derived xenografts (PDX) – where the stromal mouse DNA and mRNA component can be bioinformatically excluded “de-moused” (Guo et al., 2019) – offers a final point of independent validation. In this regard – CrownBio’s public database of multiomic characterized ∼2500 PDXs presents a good resource for final verification of homozygous deletions in specific genes.

### Recurrent homozygous deletion of the 3p25 locus center on *VGLL4* and *ATG7* but not ***VHL***

Having qualified the difficulties in identification of homozygous deletion in TCGA data collected on bulk tumors – let us consider the question of what genes are the “target” (Beroukhim et al., 2007) of the 3p25 homozygous deletion.

Initial analysis of “Deep Deletions” of the 3p25 region in the combined TCGA dataset; was performed using cBioPortal OncoPrint (summarized in **Table I**). *VGLL4* has the most numerous tumors with “Deep Deletions” followed closely by *ATG7* with a much smaller number of cases of *VHL* Deep Deletions - but as noted above the term “Deep Deletions” is not limited to homozygous deletions; instead, it is a catch all that can include actual bi-allelic, true homozygous deletions that result in complete elimination of a gene but can also include heterozygous deletions or tumors with whole genome duplications where three out of four copies of a gene have been lost. Hence this OncoPrint data lone cannot be taken as evidence of homozygous deletion. In fact – many of the cases identified as “HOMDELETED” for VHL also have VHL mutations – which would not be possible if the gene was bi-allelically (homozygous) deleted. ASCAT and ploidy corrected CNV deletion data from Sanger’s cosmic database still show a substantial number of homozygous deleted cases for *VGLL4* and *ATG7* – the number drops to only a single case (out of ∼10,000) for *VHL* when corrected for focality (**Table I**).

A detailed inspection of the segment CNV data in IGV indicates that focal low CNV regions are clustered around the partially overlapping gens *ATG7* and *VGLL4* but not in the more obvious TSG candidate – *VHL* (**Figure 1**). In fact – no focal homozygous deletions in the segmentation data are evident at the *VHL* locus (**Figure 1**). This is confirmed by GISTIC analysis (**Figure 4**) – a statistical method which aims to put a statistical quantification on focality and recurrence to identify the most likely driver of a particular CNV altered locus (Beroukhim et al., 2007). GISTIC analysis of the CNV data of the ∼10,000 tumors in TCGA indicates that *VHL* is not statistically significantly focally deleted across the entire ∼10,000 tumor dataset – and is 7.08 Mb away from the nearest statistically significant peak of focal deletion (*VGLL4/ATG7*). *VHL* is in a region of statistically significantly deletion in kidney RCC but, even in this cancer where VHL is a known TSG, *VHL* is not within the focal statistical peak region of deletion which instead covers *ATG7* and *VGLL4* (**Figure 4**). Analysis of CrownBio’s PDX data which allows unconfounded copy number calls due to bioinformatic de-convolution of stromal admixture (“de-mousing”) – shows that not a single case of *VHL*-homozygous deleted PDX could be identified (**Table I**; **Figure 5**) – but many cases with homozygous deletion in *VGLL4* and *ATG7* (**Table I**; **Figure 6**).

**Figure 4:**
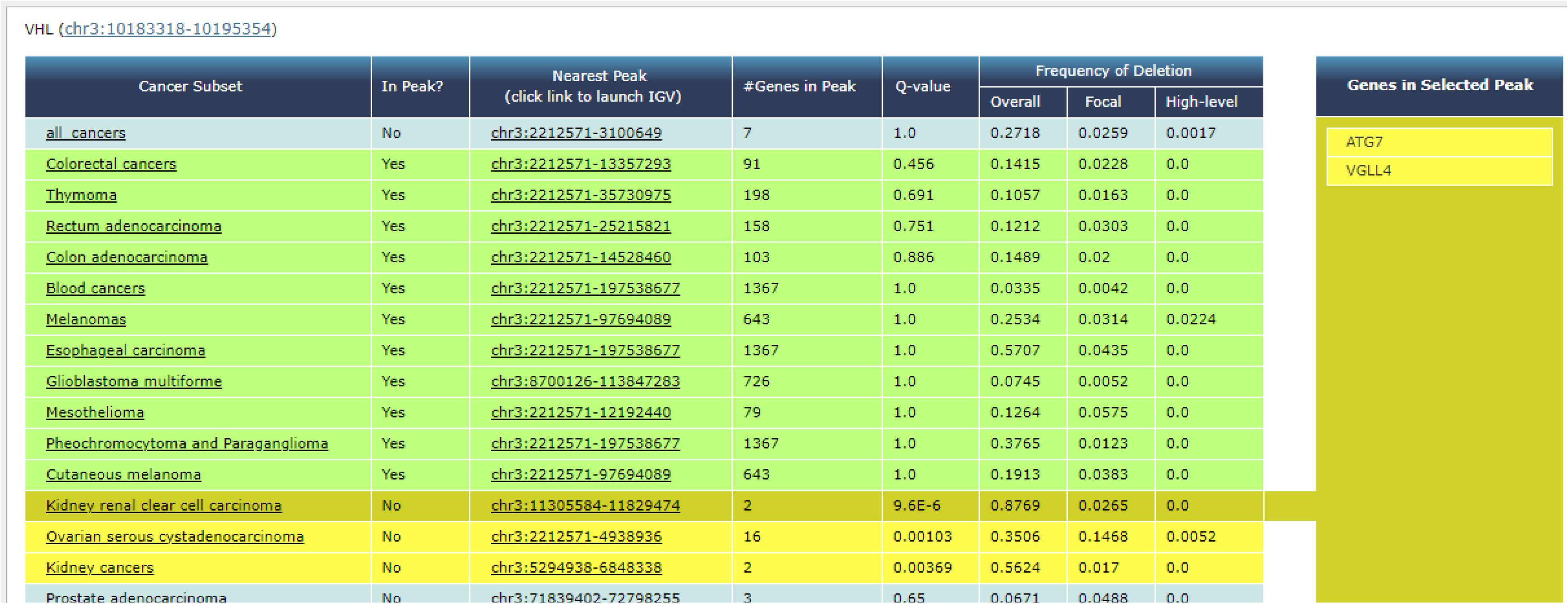
GISTIC analysis indicates that *VHL* is not focally deleted and is not within a focal peak of deletion even in RCC. The *VHL* gene was interrogated for GISTIC analyses pre-calculated and available on the Broad’s TCGA portal {https://portals.broadinstitute.org/tcga/home;(Beroukhim et al., 2010)} using the stddata 2015_04_02 TCGA/GDAC tumor sample set which represents the most up to date analysis of the dataset. Statistical analysis and interpretation are provided on the website. The analysis indicates that *VHL* is not significantly focally deleted across the entire dataset of ∼10,000 tumors (Q value > 0.05) and is not located within a focal peak region of deletion (“In Peak?” column showing “No” for the “all cancer” row). *VHL* is significantly focally deleted in 2 of 33 independent cancer types which are Ovarian serous carcinoma and RCC (Q-value <0.00001). Yet even in these two cancer types *VHL* is not in the peak of significance (“In Peak?” column showing “No” for the “Kidney cancers”, “Ovarian serous carcinoma” and Kidney “Clear Cell Carcinoma” rows). *VHL* is located 7 Mb away from the nearest peak region of deletion which is *VGLL4/ATG7*.

**Figure 5:**
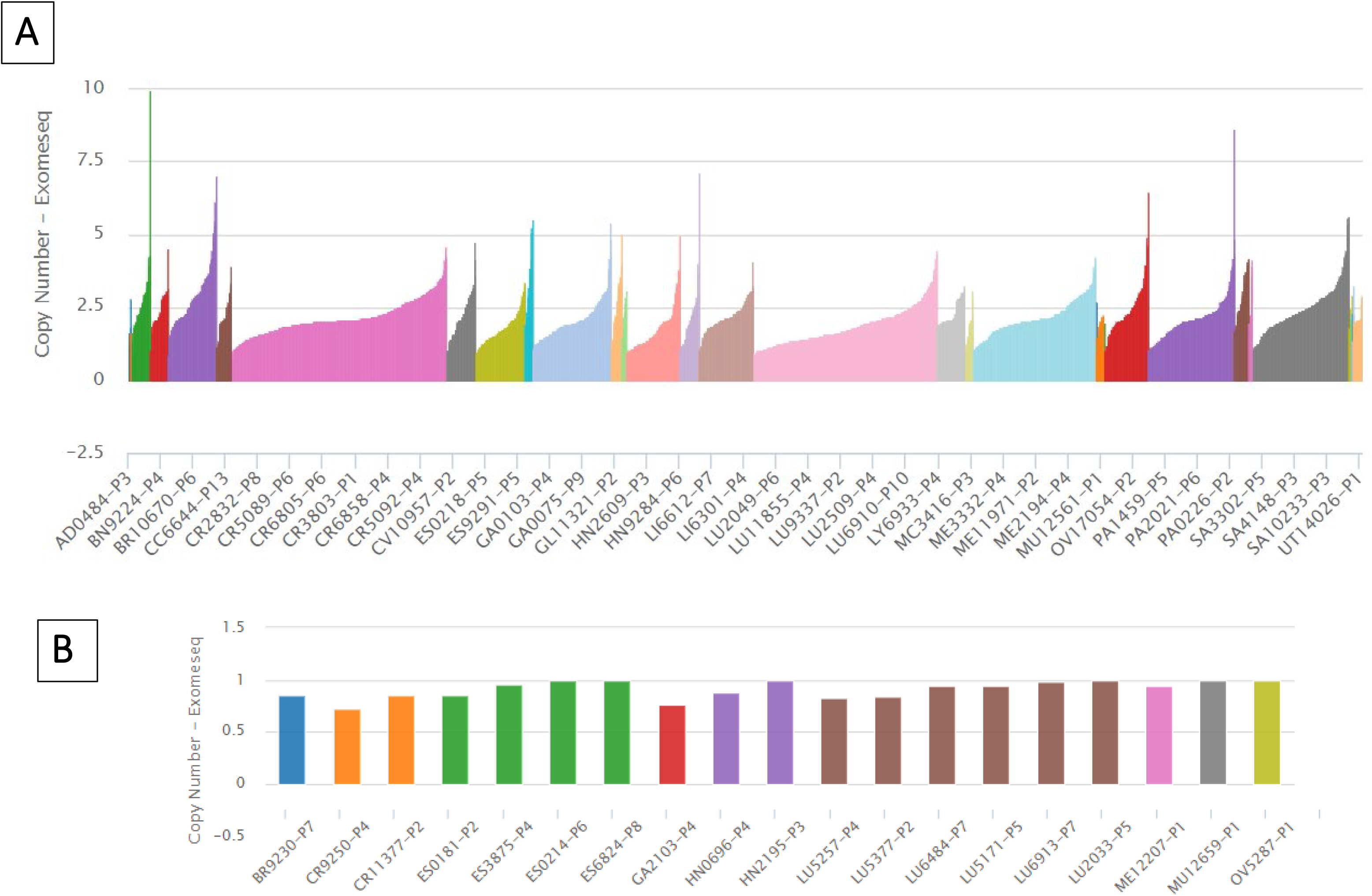
No PDX with *VHL* homozygous deletion in the CrownBio collection. CrownBio’s HuBase™ (https://www.crownbio.com/databases/hubase) of ∼2500 PDXs with de-moused Exome seq data was interrogated with *VHL* as a query for CNV. Absolute genomic copy numbers shown for PDXs of various cancer types – color-coded by cancer type. Y-axis shows genomic copy number for VHL - N = 2 is euploidy; N = 1 is a mono-allelic or heterozygous deletion; N = 0 represents bi-allelic or homozygous deletions. **Panel A**: the entire collection of ∼2500 cases. **Panel B**: subset of cases with copy number less than 1 – note none are near zero – indicating no PDX with homozygous deletion of *VHL*.

**Figure 6:**
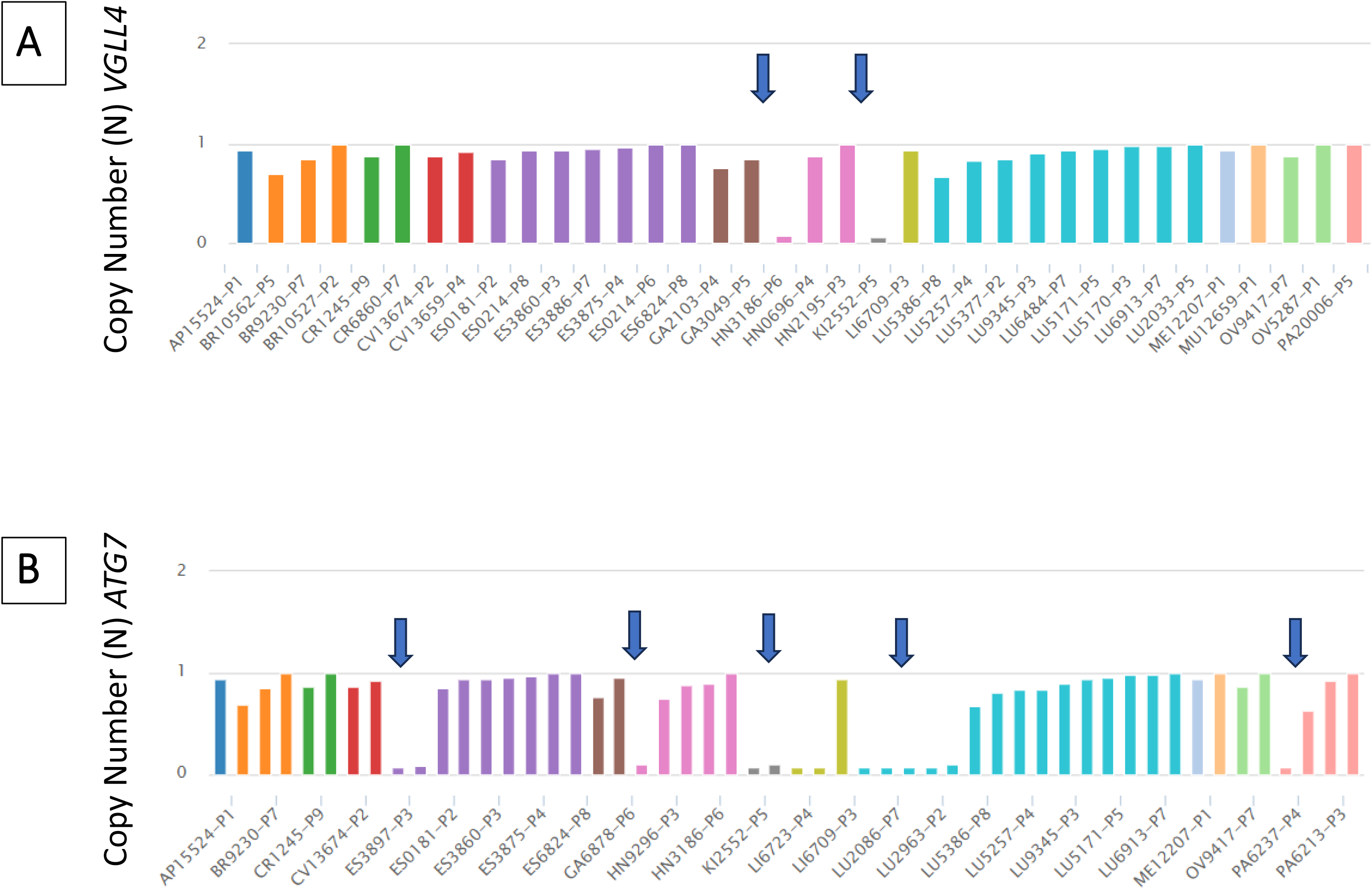
Recurrent homozygous deletions of *VGLL4* and *ATG7* in the CrownBio PDX collection. CrownBio’s HuBase™ was interrogated with *VGLL4 (***Panel A**) and *ATG7 (***Panel B**) as queries for CNV. Absolute genomic copy numbers derived from de-moused whole exome sequencing data for PDXs of various cancer types – color-coded by cancer type. Y-axis shows absolute genomic copy number- N = 2 is euploid; N = 1 is a mono-allelic or heterozygous deletion; N = 0 represents bi-allelic or homozygous deletions. Panel A: Subset of cases with copy number less than 1 for VGLL4 with blue arrows indicating homozygous deletions of *VGLL4* (Panel A) and *ATG7* (Panel B). The cancer types affect closely match those identified in TCGA data – in squamous cell carcinoma especially lung. To highlight a specific case: a kidney clear cell carcinoma PDX KI2552 has a homozygous deletion spanning *VGLL4* and *ATG7*.

A final dataset to evaluate the TSG candidate of the 3p25 homozygous deletion is the CRISPR and RNAi Dependency data from DepMap (Tsherniak et al., 2017). DepMap RNAi and CRISPR ablation of *VHL* is cytotoxic to most cell lines (**Figure 7A**; **Table I**) with the gene being meeting the “essential” threshold in the vast majority of DeMap cell lines. It is only in RCC mutant that VHL CRISPR dependency Chronos score is around 0 (fitness neutral) though even some *VHL*-mutant RCC cell lines show strongly cytotoxic scores. In contrast – extinction of *VGLL4* (and to a lesser extent – ATG7) strongly promotes proliferation and shows a CRISPR profile similar to well-established TSGs (**Figure 8**).

**Figure 7.**
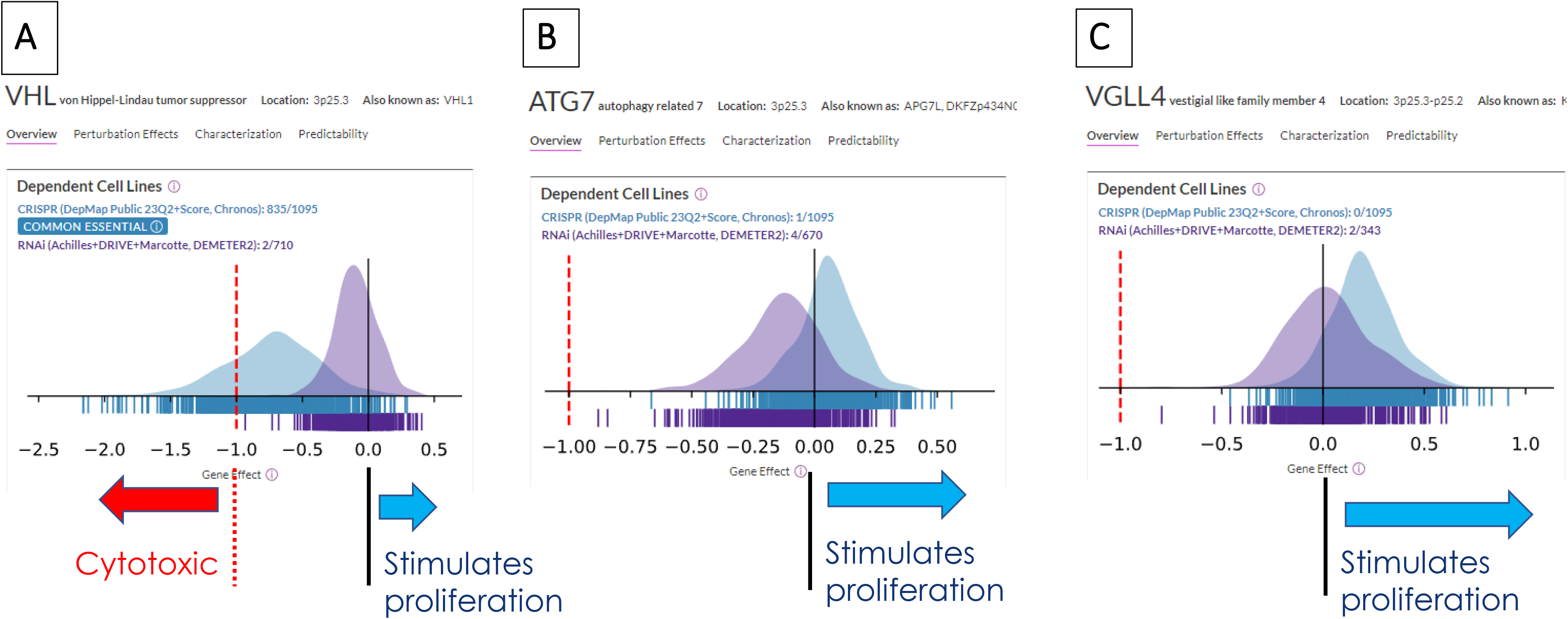
DepMap CRISPR dependency data for 3p25 candidate target TSG indicates that *VHL* is, unexpectedly, an essential gene in many cancer cell lines. The Broad’s DepMap portal (https://depmap.org/portal/) was interrogated for CRISPR and RNAi dependencies of 3p25 candidate genes *VHL*, *VGLL4* and *ATG7* to determine whether their homozygous deletion would be a strong oncogenic driver event. Summary of CRISPR Chronos scores for 23Q2 data are shown for *VHL* (**Panel A**), *ATG7* (**Panel B**) and *VGLL4* (**Panel C**) – negative values indicate that CRISPR ablation is detrimental whilst positive values indicate that CRISPR ablation increases fitness and stimulates cancer cell proliferation. Chronos values below −1 indicate essential genes which is indicated by a red dashed line. Each bar represent one cell line. CRISPR cell line dependency data for *VHL* unexpectedly recapitulate those of core housekeeping genes (e.g. *PCNA*, *CDK1* and *POLA2* Figure 2) rather than those of typical TSG such as *TP53 or CDKN2A*. *VHL* CRISPR knockout is detrimental in the vast majority of cancer cell lines and meets the essential threshold in 835 cells lines out of 1095. CRISPR data for ATG7 on balance are skewed in the positive direction - indicating that *ATG7* ablation is neutral or weakly stimulates cell proliferation across most cancer cell lines, providing support for *ATG7* homozygous deletion as a driver event. *VGLL4* CRISPR data most closely align with those of a classic TSG – *VGLL4* knockout results in increased fitness and proliferation in most cell lines and the median Chronos score is strongly positive.

**Figure 8:**
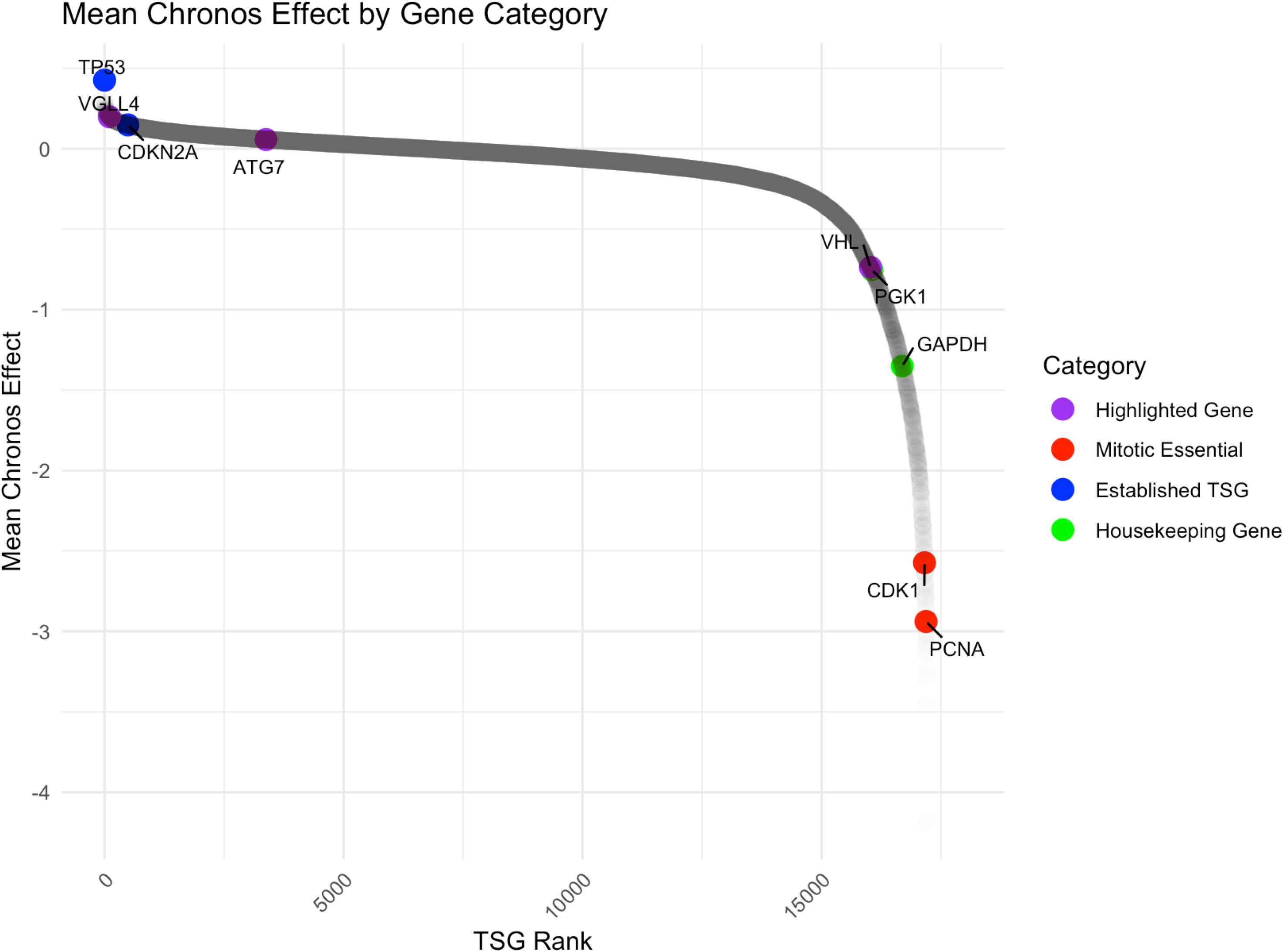
*VGLL4* and *ATG7* rank highly as tumor-suppressors in the DepMap while *VHL* does not. DepMap (23Q2) data were interrogated for CRISPR Dependency scores. Mean Chronos Effect scores were calculated for each gene across all available cell-lines. TSG ranks were derived by ranking genes by Mean Chronos Effect, where a larger positive Mean Chronos Effect indicates that gene knockout leads to increased proliferation and a negative score indicates determinantal effects. To contextualize – card-carrying TSG (blue) TP53 and CDKN2A are ranked #5 and #529 whilst established metabolic housekeeping genes (green) like *GAPDH* and *PGK1* are ranked #17394 and #16740 and essential mitotic genes (red) such as CDK1 and PCNA, are at the very bottom of the TSG scale.

Taken together – the data offer support for both *VGLL4* and *ATG7* as the TSG target of the *3p25* homozygous deletion and warrant a more detailed look at the tumor suppressive function and cancer types most affected.

### *VGLL4*-homozygous deletion as a driver event in Carcinomas

GISTIC analysis of *VGLL4* indicates strong statistical significance for focal deletion in the whole TCGA ∼10,000 tumor dataset (reflecting 3p deletions) though *VGLL4* is not in the minimal common region of deletion in this pan-cancer group (**Figure 9**). However, in Lung squamous cell carcinoma and carcinomas as a group (tumors of epithelial origin) *VGLL4* is strongly significance for focal deletion and is the only gene in the peak of minimum common deletion. In RCC, Esophageal carcinoma and Cholangiocarcinoma – both *VGLL4* and *ATG7* are in the peak of significance. The cancer-centric GISTIC analysis indicates that 3p25 is among the most statistically significant loci of focal deletion in lung squamous cell carcinoma (**Figure 10A**).

**Figure 9:**
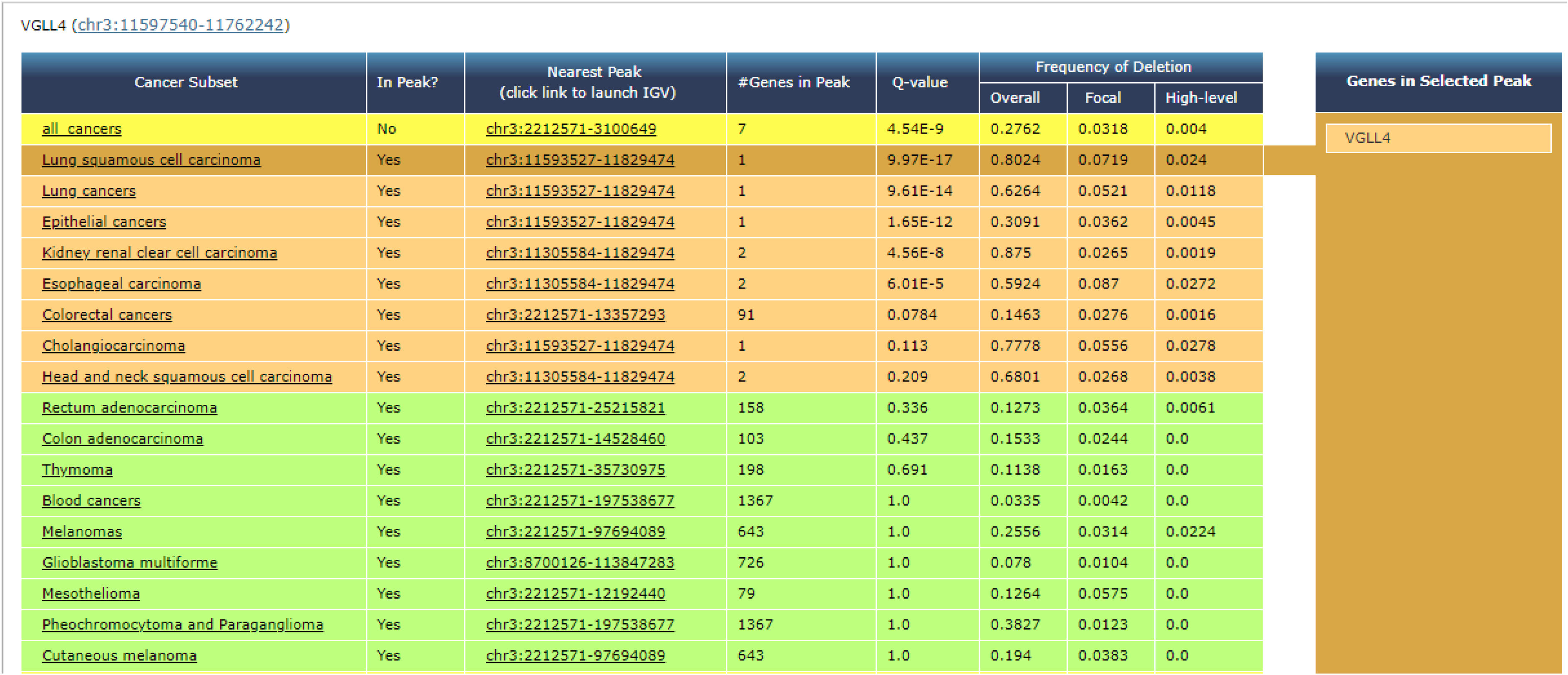
GISITC gene-centered analysis indicates statistically significant focal deletion of *VGLL4* in carcinomas. The *VGLL4* gene was interrogated for GISTIC gene-centered pre-calculated analyses of the Broad’s TCGA portal using the stddata 2015_04_02 TCGA/GDAC tumor sample set. *VGLL4* is significantly focally deleted across the entire dataset but is not located within a focal peak region of deletion (“In Peak?” column showing “No” for the “all cancer” row). *VGLL4* is significantly focally deleted in several independent cancer types and in epithelial cancers (Carcinomas) as a group. *VGLL4* is within the region of minimum common deletion and is the sole gene in the peak in Lung Squamous Cell Carcinoma and carcinomas as a group. In RCC and Esophageal carcinomas *VGLL4* is one of two genes in the peak – with *ATG7* being the other. Together – these data provide strong statistical support for VGLL4 as the target of the 3p25 deletion.

**Figure 10:**
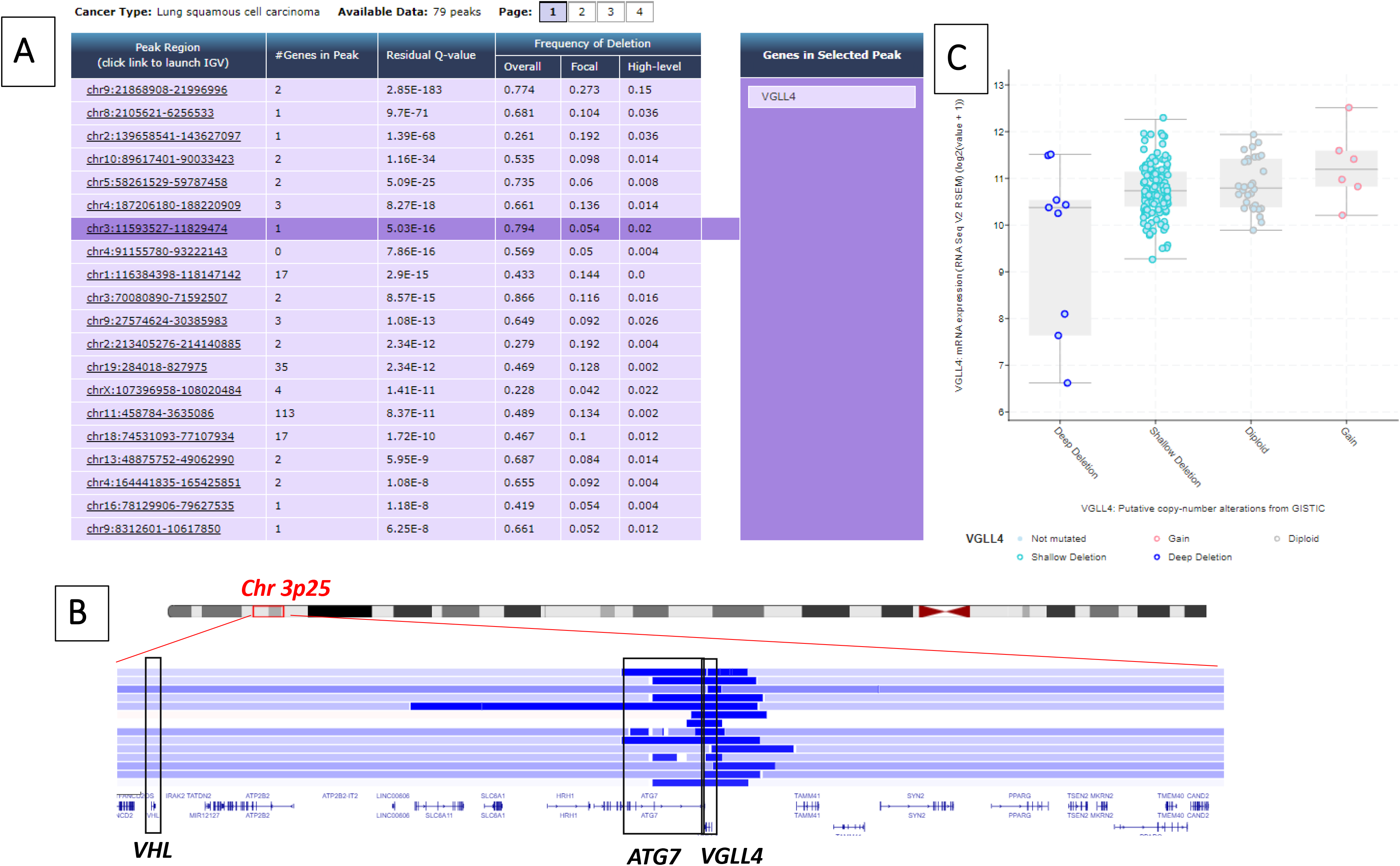
Recurrent focal homozygous deletions centered on *VGLL4* in lung squamous cell carcinoma. Lung squamous cell carcinoma was interrogated for GISTIC cancer-centered pre-calculated analyses of the Broad’s TCGA portal using the stddata 2015_04_02 TCGA/GDAC tumor sample set. *VGLL4* was the 7^th^ most significantly deleted gene – both focal; recurrent and the only gene in the peak of minimum region of common deletion (**Panel A**). A detailed look at the segmented SNP 6.0 data indicates homozygous deletions centered on *VGLL4* rather than *AGT7 (***Panel B**). Cases with Deep Deletion of *VGLL4* show dramatically (several Log2-fold0 lower mRNA expression (**Panel C**) reinforcing that in the case of *VGLL4,* instances of Deep Deletions indeed correspond to homozygous deletions.

When considering only homozygous deletions the 3p25 locus is the third most frequently deleted locus in Lung squamous cell carcinoma – after 9p21 (*CDKN2A*) and 10q21 (*PTEN*). A detailed segmentation of analysis of SNP 6.0 data in lung squamous cell carcinoma with IGV (**Figure 10B**) nicely illustrates the pattern of homozygous deletion skewing towards *VGLL4* versus *ATG7*.

Further analysis of mRNA expression in the RNA seq TCGA data complements and reinforces the CNV data – showing robust reduction in mRNA expression of several Log2-fold in tumors with high-confidence homozygous deletion of *VGLL4* (**Figure 11**). Cases with “deep deletions” (dark blue circles) show several Log2-fold lower mRNA expression and genuine tumors with homozygous deletions of *VGLL4*. Shallow deletion case with very low mRNA expression, pointed out in light blue arrows, may also represented homozygous deleted tumors with high stromal admixture. While *VGLL4* is expressed at robust levels in most genomically intact tumors – some cancer types such as Hepatocellular carcinoma, Pheochromocytoma, and Seminoma show group-level decreases in *VGLL4* (**Figure 11**).

**Figure 11:**
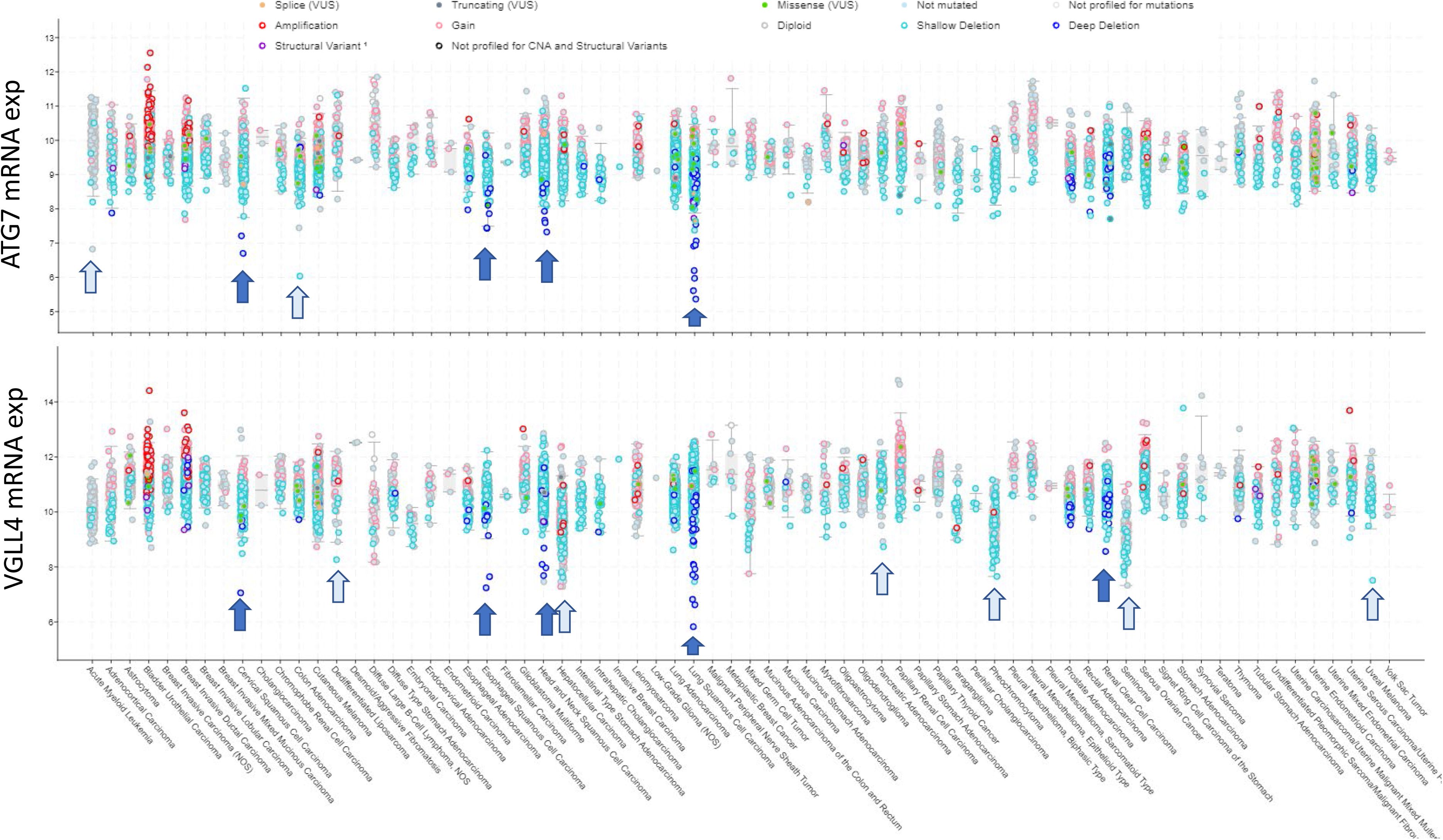
Reduced mRNA expression of *VGLL4* and *ATG7* through homozygous deletions and lineage in TCGA. VGLL4 (lower panel) and ATG7 (upper panel) were queried in cBioPortal’s assembled TCGA combined tumor dataset. Expression (mRNA seq Log2 on y-axis) was plotted versus specific disease (x-axis) with each dot representing a tumor case with color coding reflecting point mutation or CNV calls as indicated. Cases with deep deletions (dark blue circles) frequently show dramatically lower mRNA expression (several log2-fold) and reflect genuine homozygous deletions (marked by blue arrows). This concordance of mRNA and copy number is reflected in the recurrent deletions in Lung squamous cell carcinoma. Shallow deletions (heterozygous) with very low mRNA expression are pointed out in (light blue arrows). Lineages with very high frequency heterozygous deletion (shallow deletion) and dramatically lower *VGLL4* expression include Hepatocellular carcinoma; Pheochromocytoma and Seminoma.

Mechanistically – the oncogenic impetus of the homozygous deletion of VGLL4 has a straightforward explanation: VGLL4 is THE main negative regulator of the YAP/TAZ-TEAD transcriptional complex and VGLL4 loss is expected to result in hyperactivation of transcriptional targets linked to proliferation, de-differentiation, and survival (**Figure 12**). That *VGLL4*-homozygous deletion truly is a major driver event finds extensive support in bioinformatics databases and the literature. DepMap CRISPR KO data make a very strong case for homozygous deletion of VGLL4 as a major driver event: CRISPR KO of VGLL4 promotes fitness and stimulates proliferation in the vast majority of cell lines (**Table I**; **Figure 7C**). Indeed, VGLL4 and is amongst the top 150 genes in the DepMap whereby CRISPR KO induces proliferation in the DepMap, along with well-known TSGs like PTEN and TP53 (Figure if needed). This is complemented by extensive single-cell lines shRNA studies {e.g. (Zhang et al., 2014)}. Few functional validation studies with genetically engineered mouse models (GEMM) for homozygous deletion of *VGLL4* as a driver have been conducted (Cai et al., 2022). Indeed GEMM studies have mostly focused on the role of VGLL4 in development (Sheldon et al., 2022) rather than cancer. It is only very recently because of the recognition of VGLL4’s importance as a negative regulator of the YAP/TEAD oncogenic transcriptional complex (**Figure 12**) that dedicated GEMM studies with cancer have been undertaken. In a landmark study Cai et al. demonstrated that homozygous deletion of *Vgll4* strongly stimulated YAP/TEAD transcriptional activity and cell proliferation and dramatically enhanced intrahepatic cholangiocarcinoma (ICC) formation in combination with *Nf2* 11/6/2023 7:07:00 AM. It is noteworthy that while Cai et al. keenly appreciate the importance of their finding vis-a-vis the Hippo pathway in context – they appear unaware of the homozygous deletion of *VGLL4* noted in the present publication. More broadly while the heterozygous deletion of *VGLL4* is well known to the community due to high frequency 3p chromosome arm deletions (He et al., 2021) – even publications that have made detailed investigations into genetic alterations of the Hippo pathway did not discuss VGLL4 homozygous deletions (He et al., 2021).

**Figure 12:**
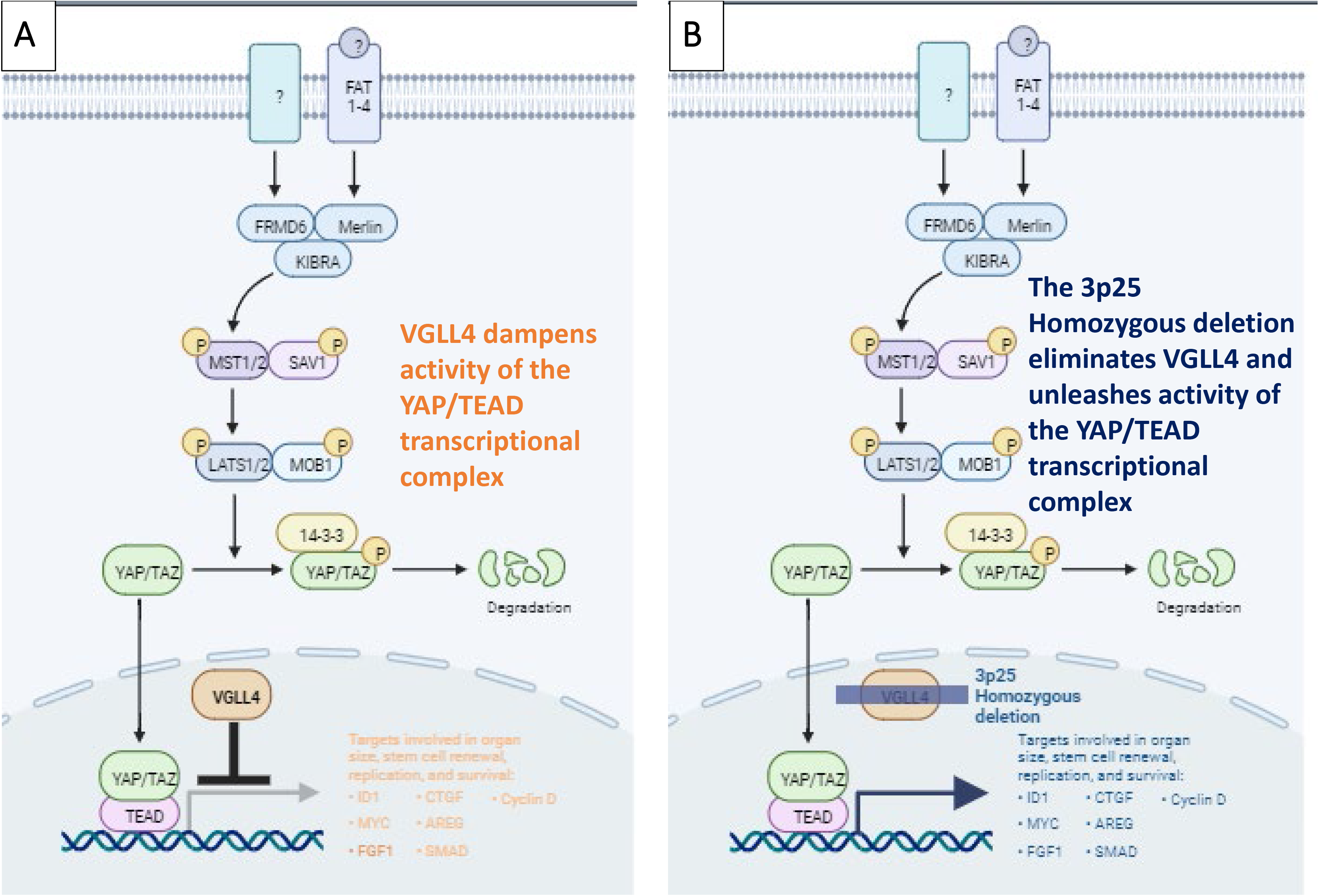
The 3p25 Homozygous deletion eliminates VGLL4 and unleashes oncogenic activity of the YAP/TEAD complex. **Panel A:** VGLL4 and YAP/TAZ compete for the same binding-domain on TEAD transcription factors (Vaudin et al., 1999) with VGLL4 acting as a direct repressor of YAP-dependent TEAD transcriptional activity {(Koontz et al., 2013);(Mickle et al., 2021);(Jiang et al., 2015)}. **Panel B:** homozygous deletion of 3p25 locus eliminates VGLL4, de-repressing the transcriptional activity of the YAP/TEAD complex and oncogenesis through transcriptional target genes that promoting proliferation, de-differentiation and survival.

Taken together – the genomic data in TCGA and the functional mechanistic data from both cell lines and mice – make a very strong case for *VGLL4* homozygous deletion as a key oncogenic driver event.

### *ATG7*-homozygous deletion as a driver event in RCC

GISTIC analysis of the 3p25 locus generally provides stronger support for *VGLL4* as compared to *ATG7* as the target of the deletion – except in RCC where *ATG7* is included in both the focal region of deletion and in the peak of highest significance (**Figure 4**). In addition – *ATG7* homozygous deletion is evident in multiple PDXs (**Figure 6**). There also significant number of ATG7 mutations that although not labeled as “hotspot” are likely to be damaging – generating loss-of-function which given the frequent arm-level heterozygous deletions of chromosome 3p would effectively inactivate *ATG7* in cancer cells (**Table I**). Interestingly – this exact scenario was documented in familial cholangiocarcinoma where inherited *ATG7*-hypomorphic mutations resulted in cholangiocarcinoma with null alleles due to heterozygous deletion of the remaining WT allele (Greer et al., 2022). GEMM studies indicate that germline *Atg7* knockout mice die shortly after birth but that mosaic knockout mice develop hepatic adenomas (Takamura et al., 2011). CRISPR data generally indicate that ATG7 ablation stimulates proliferation, but the effect is overall much weaker than with VGLL4 (**Table I**; **Figure 7B**). But this might be expected from the mechanism of tumor suppression – which would not be manifest in cell culture under abundant nutrient conditions. It has been proposed that ATG7 limits proliferation in response to nutrient limitation through a p53 dependent mechanism (Lee et al., 2012). Since emerging malignancies rapidly outgrow their nutrient supply – ATG7 likely limits proliferation and its inactivation either by point mutation coupled with a heterozygous deletion or by bi-allelic (homozygous deletion).

Previous studies had noted that ATG7 may act as a tumor suppressor and one study even identified an ATG7-null cell line, H1650 (Mandelbaum et al., 2015), as a useful tool for studying autophagy. Yet no previous study reported that ATG7 is recurrently homozygous deleted in diverse human tumors and that this deletion might be exploited for precision oncology targeting.

## Discussion

What is the target tumor suppressor gene of the 3p25 homozygous deletion and by what mechanisms does it drive tumor progression? And importantly – what are the therapeutic implications of the answers to this question?

The most obvious candidate is *VHL* (negative regulator of HIF) but as this inquiry found, *VGLL4* and *ATG7* are the actual targets. Analysis of the minimal common region of deletion places *VHL* outside the peak of significance – indeed despite being frequently inactivated by combination of heterozygous deletion + point mutation – *VHL* homozygous deletions are rare even in kidney RCC. The CRISPR scores for VHL would imply that it might serve as an oncology drug target though the organismal toxicity of VHL inhibitors would likely be significant. This leaves *VGLL4* as the obvious candidate – an attractive candidate given its importance as a negative regulator of the pro-oncogenic YAP/TEAD transcriptional complex (**Figure 12**). In squamous cell carcinomas *VGLL4* does indeed reach the highest level of GISTIC statistical significance.

DepMap CRISPR data indicate that *VGLL4* ablation strongly increases proliferation in diverse cancer cell lines (**Figure 7B**) which together with GEMM data (Cai et al., 2022) strongly supports *VGLL4*-deletion as a genuine driver event. Therapeutically – the Hippo pathway has recently become actionable with palmitic acid site TEAD inhibitors {Vivace; (Tang et al., 2021)} and YAP/TAZ inhibitors having entered the clinic {Novartis; (Furet et al., 2022)}. It stands to reason that *VGLL4*-homozygous deleted tumors would be good candidates for such therapies (**Figure 12**). The homozygous deletion of *VGLL4* is highly recurrent in several common cancer types such as esophageal and lung squamous cell carcinoma – translated into actionable treatments for perhaps 10,000 or patients per year in the U.S.

Because the genes encoding *VGLL4* and its immediate neighbor *ATG7* are on opposite DNA strands and partially overlap, deletion of both genes is quite common and discerning, which is the “driver” is rather complex. While the GISTIC scores in most cancer types provided stronger support for *VGLL4* in most cancer, in selected types such as RCC, *ATG7* is in the peak of significance.

*ATG7* is well known for its central role in autophagy (Collier et al., 2021) and the immediate thought would be that deletion of such as housekeeping gene constitutes a passenger co-deletion rather than an actual TSG target of the 3p25 deletion. Yet the GISTIC peak clearly centers on *ATG7* in RCC and this data is strengthened by analysis of PDXs banks where homozygous deletion calls are less confounded (**Table I**; **Figure 6**). A careful look at the mechanistic role of ATG7 in autophagy indicates that it may well act as a tumor suppressor and that its deletion is a driver event: ATG7 is activated under nutrient-limiting conditions and leads to growth arrest (Lee et al., 2012) somewhat akin to AMPK/LKB1 (Alessi et al., 2006). It is tempting to speculate that *ATG7*-homozygous deletion acts akin to LKB1 loss-of-function, allowing continuous cancer cell proliferation even under nutrient limiting conditions. Compelling evidence for a direct classic Knudson 2-hit tumor suppressor role of ATG7 comes from ultra-rare familial cholangiocarcinoma – where a loss-of-function *ATG7* point mutation segregates with cholangiocarcinoma incidence and where the WT allele is lost by 3p arm deletion, rendering such tumors functional ATG7-null (Greer et al., 2022).

Tumors with *ATG7*-homozygous deletions present a unique synthetic lethal opportunity that has not been explored – largely because this genetic event is not present in established cancer cell lines such as those found in the major cell line banks like DepMap. How might ATG7-homozygous deletions be targeted therapeutically? Normal cells activate *ATG7* to limit cell proliferation in response to nutrient and energy scarcity. It stands to reason that ATG7-null cells would continue to proliferate – to the point of starvation – even in states of nutrient deprivation. This suggests that ATG7-null cancers may be susceptible to metabolic targeted therapies.

In sum – the homozygous deletion of *VGLL4* and *ATG7* represent heretofore unrecognized recurrent genomic drivers in diverse cancers and potential targetable vulnerabilities in precision oncology. Because the significance of these deletions was not previously recognized – *VGLL4* and *ATG7* are not currently included even in the most extended targeted cancer sequencing panels such as the TEMPUS xT, which includes over 600 genes (https://www.tempus.com/wp-content/uploads/2023/06/Tempus-xT_Gene-Panel.pdf). Our findings urgently call for the inclusion of ATG7 and VGLL4 in more advanced sequencing panels.

